# Single-cell immune landscape of human recurrent spontaneous abortion

**DOI:** 10.1101/2020.09.16.300939

**Authors:** Feiyang Wang, Wentong Jia, Mengjie Fan, Zhilang Li, Yongjie Liu, Yeling Ma, Xuan Shao, Yu-xia Li, Rong Li, Qiang Tu, Yan-Ling Wang

## Abstract

Successful pregnancy in placental mammals substantially depends on the establishment of maternal immune tolerance to the semi-allogenic fetus. Disorders in this process are tightly associated with adverse pregnancy outcomes including recurrent spontaneous abortion (RSA). However, an in-depth understanding of the disorders from the aspect of systematic and decidual immune environment in RSA remains largely lacking. In this study, we utilized single-cell RNA-sequencing to comparably analyze the cellular and molecular signatures of decidual and peripheral leukocytes in normal and RSA pregnancies at the early stage of gestation. Integrative analysis identified 22 distinct cell clusters in total, and a dramatic difference in leukocyte subsets and molecular properties in RSA cases was revealed. Specifically, the cytotoxic properties of CD8T effector, NK, and MAIT cells in peripheral blood indicated apparently enhanced immune inflammatory status, and the subpopulation proportions and ligand-receptor interactions of the decidual leukocyte subsets demonstrated preferential immune activation in RSA patients. The molecular features, spatial distribution and the developmental trajectories of five decidual NK (dNK) subsets were illustrated. The proportion of a dNK subset responsible for fetal protection was reduced, while the ratio of another dNK subset with cytotoxic and immune-active signature was significantly increased. Notably, a unique pro-inflammatory CD56+CD16+ dNK subpopulation was substantially accumulated in RSA decidua. These findings reveal a comprehensive cellular and molecular atlas of decidual and peripheral leukocytes in human early pregnancy, which provides an in-depth insight into the immune pathogenesis for early pregnancy loss.

## INTRODUCTION

Reproductive success in placental mammals substantially depends on the establishment of maternal immune tolerance to the semi-allogenic fetus during pregnancy (Ander et al., 2019; Erlebacher, 2013). In humans, failures in immune tolerance are tightly associated with various adverse pregnancy outcomes, mainly including recurrent spontaneous abortion (RSA), preeclampsia, and stillbirth (Deshmukh and Way, 2019; Fisher, 2015; Kheshtchin et al., 2010)

The key of fine-tuned immune tolerance at the feto-maternal interface is the well-coordinated crosstalk among uterine mucosal immune cells and extraembryonic placental cells. The maternal immune cells populating the uterine mucosa include decidual NK (dNK) cells (∼50-70%), macrophages (∼20%), T cells (∼10-20%) and a small amount of DC and B cells. The numerically dominant dNK cells are characterized by low cytotoxicity and strong cytokine producing capacity, which is divergent from peripheral NK (pNK) cells (Wang et al., 2014). Mechanistically, the inhibitory receptors LILRB2 and KIR2DL4 on dNK cells recognize HLA-G on the surface of trophoblast cells, leading to the suppressed cytotoxic capacity of dNK cells (Gamliel et al., 2018a; Parham and Moffett, 2013). Moreover, the biased differentiation of CD4+ T cells towards Th2 and Treg has been confirmed vital to the tolerance of fetal cells in decidua (Wang et al., 2010; Yang et al., 2008).

Murine decidual immune cells, especially innate lymphocytes, have been characterized as multiple subsets that present dynamical lineage differentiation and crosstalk properties during pregnancy (PrabhuDas et al., 2015). However, the corresponding systematic study on human decidua is largely lacking, which hampers the in-depth understanding of the physio-pathological regulation of human pregnancy. A recent paradigm shifting study by Moffett and colleagues utilizing single-cell RNA-sequencing and comprehensive data analysis constructed a detailed cellular map and the elaborate cell-cell communication patterns of the human decidual-placental interface. Specifically, three major subsets of dNK cells that have distinctive immunomodulatory and chemokine profiles were identified (Huhn et al., 2020; Suryawanshi et al., 2018; Vento-Tormo et al., 2018). However, high-resolution immune landscape of dysregulated decidua in the contexts of pregnancy complications, for example in RSA, is still unknown, which greatly limit the exploration of immune-related pathogenesis for recurrent miscarriage.

In the present study, we utilized single-cell RNA-sequencing to comparably analyze the cellular and molecular signatures of decidual and peripheral leukocytes in normal and RSA pregnancies at the early stage of gestation. By integrating the gene expression properties, ligand-receptor interactions, and spatial localization patterns, the cell-type-specific communications among various subsets of leukocyte and the developmental trajectory of dNK cells were constructed. A pathological leukocyte atlas in RSA decidua and peripheral blood was illustrated. Our findings reveal a detailed molecular and cellular map of decidual and peripheral leukocytes in early pregnancy of human, and highlight the integral harmful immune responses that may lead to early pregnancy failure.

## RESULTS

### Clustering of immune cells from peripheral blood and decidual tissues in normal and RSA pregnancies

To characterize the immune cells in RSA, we applied scRNA-seq to study CD45+ cells isolated from peripheral blood and decidual tissues of three pairs of normal pregnant women and RSA patients with unknown cause. Viable CD45+ leukocytes were enriched by FACS and subjected to droplet-based single-cell RNA sequencing (scRNA-seq) using the 10x Genomics Chromium platform (Fig. S1A and B). Sequencing data from six peripheral blood samples and six decidual tissue samples were obtained from 12 scRNA-seq libraries. Following computational quality control and filtering using the Seurat package (Butler et al., 2018; Stuart et al., 2019), the final datasets containing 56,758 high-quality cells were subjected to further analysis (Fig. S1C, D and E).

We performed an unsupervised graph-based clustering to analyze scRNA-seq data in Seurat (version 3.0.3). In order to eliminate unreasonable clustering due to batch effects, we integrated the data using the reciprocal PCA function of Seurat. Overall, 22 transcriptionally distinct clusters were identified (Fig. 1A, B and C), and were further confirmed using the MNN algorithm (Haghverdi et al., 2018)(Supplementary Fig. S2). The cell identity of the 22 clusters was assigned on the basis of known marker genes and literature evidence (Vento-Tormo et al., 2018)(Suryawanshi et al., 2018) (Fig. 1D).

**Fig. 1.**
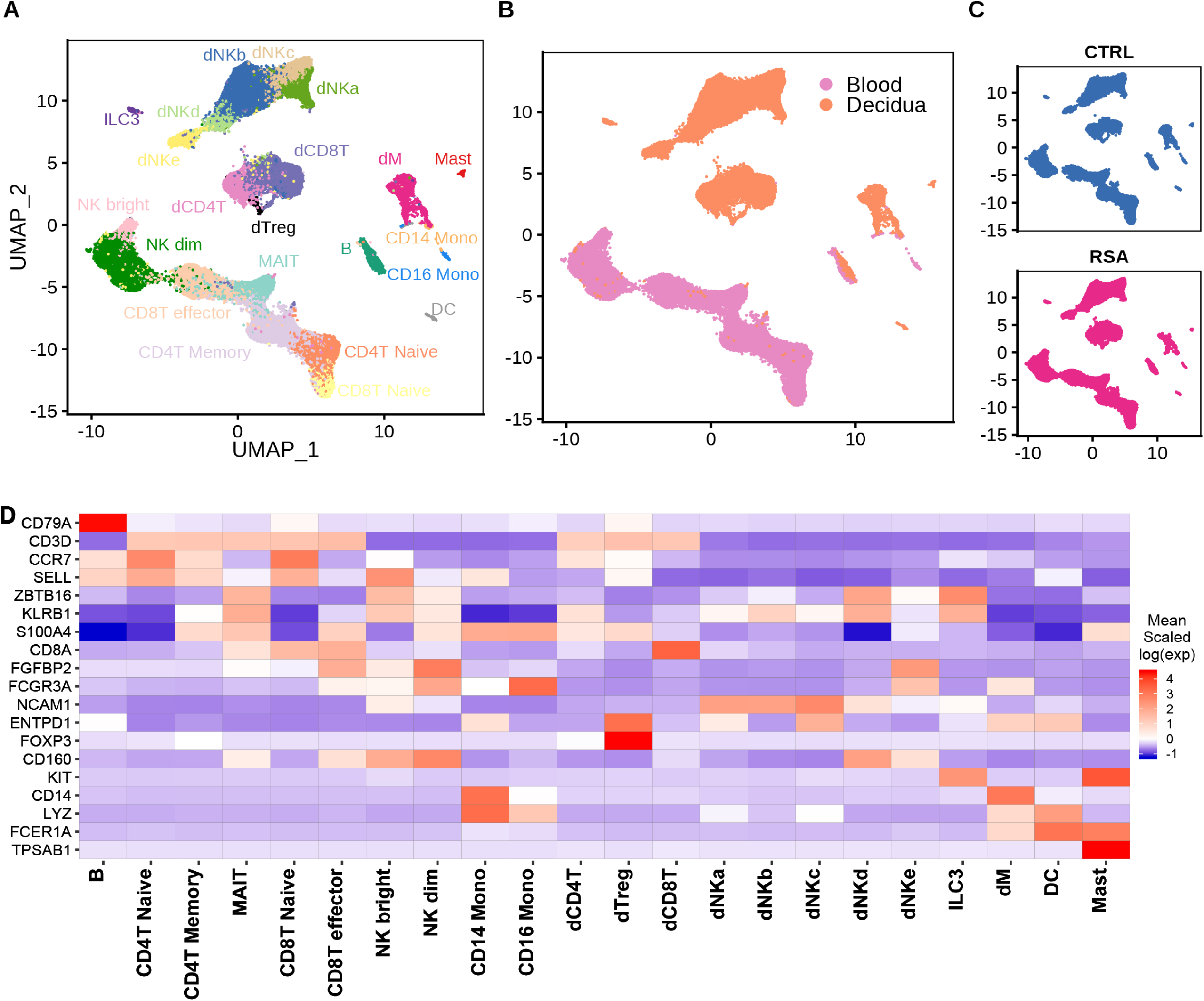
Landscape of immune cells from peripheral blood and decidual tissues in normal pregnant women and RSA patients at early pregnancy using single-cell RNA-seq. (A) UMAP plot of scRNA-seq data to show the 22 leukocyte clusters in peripheral blood and decidual tissues at early pregnancy. (B) UMAP plot showing cell populations from peripheral blood and decidual tissue. (C) UMAP visualization of cell clustering in normal and RSA pregnancies. (D) Characteristics of the 22 leukocyte clusters based on canonical marker genes expression for cell type annotation.

In peripheral blood, T cells were identified and grouped into five major clusters, including two subsets of Naïve T cells (CD4T Naive and CD8T Naive) which express the marker gene *CCR7*, mucosal-associated invariant T cells (MAIT) specifically expressing *ZBTB16*, CD4 Memory T cells (CD4T Memory) annotated by *S100A4*, and CD8 T effector cells (CD8T effector) that express marker genes *FCGR3A* and *FGFBP2* (Fig. 1D; Supplementary FigS3). Peripheral NK cells were clustered as CD56dimCD16+ NK cells (NK dim) and CD56brightCD16-NK cells (NK bright). Monocyte included CD14+ subset and CD16+ subset. In addition, a low abundance of of B cells (identified by *CD79A* expression) and DC cells (annotated by *LYZ* and *FCER1A*) were captured in peripheral blood.

In decidual tissues, the most abundant population appeared as dNK cells, which were subgrouped into five clusters, named as dNKa, dNKb, dNKc, dNkd, and dNKe. The dNKe subset presented positive expression for *FCGR3A*/CD16, which was uniquely accumulated in RSA decidual tissues. Decidual T cells were clustered as CD4+, CD8+, and a relatively small number of FOXP3+ regulatory T cells (dCD4T, dCD8T, and dTreg). Other decidual leukocyte clusters included *KIT*-expressing ILC3 cells (ILC3), *CD14*-expressing macrophage (dM), FCER1A-expressing dendric cells (DC), and TPSAB1-marked mast cells (Mast) (Fig. 1A and D).

### Cellular and molecular characteristics of the peripheral leukocytes indicate a systematic pro-inflammatory feature in RSA patients

Given T cells present a dominant proportion in peripheral leukocytes, we therefore first analyzed the properties of T cell clusters in RSA and normal pregnancies. The gene expression profiles of the five peripheral T cells subsets were indistinguishable between RSA and normal pregnancies (Fig. 2A). However, the proportion of each T cell cluster exhibited evident alterations in RSA patients, presenting by reduced CD4 T Naïve, CD8 T Naïve and CD4 T Memory populations, and augmented CD8 T effector and MAIT populations (Fig. 2B). In addition, transcription of inflammatory factors, such as *IRF1, RORA* in MAIT cells and *IFNG* in CD8 T effector cells and what was remarkably increased in RSA cases (Fig. 2C and D).

**Fig. 2.**
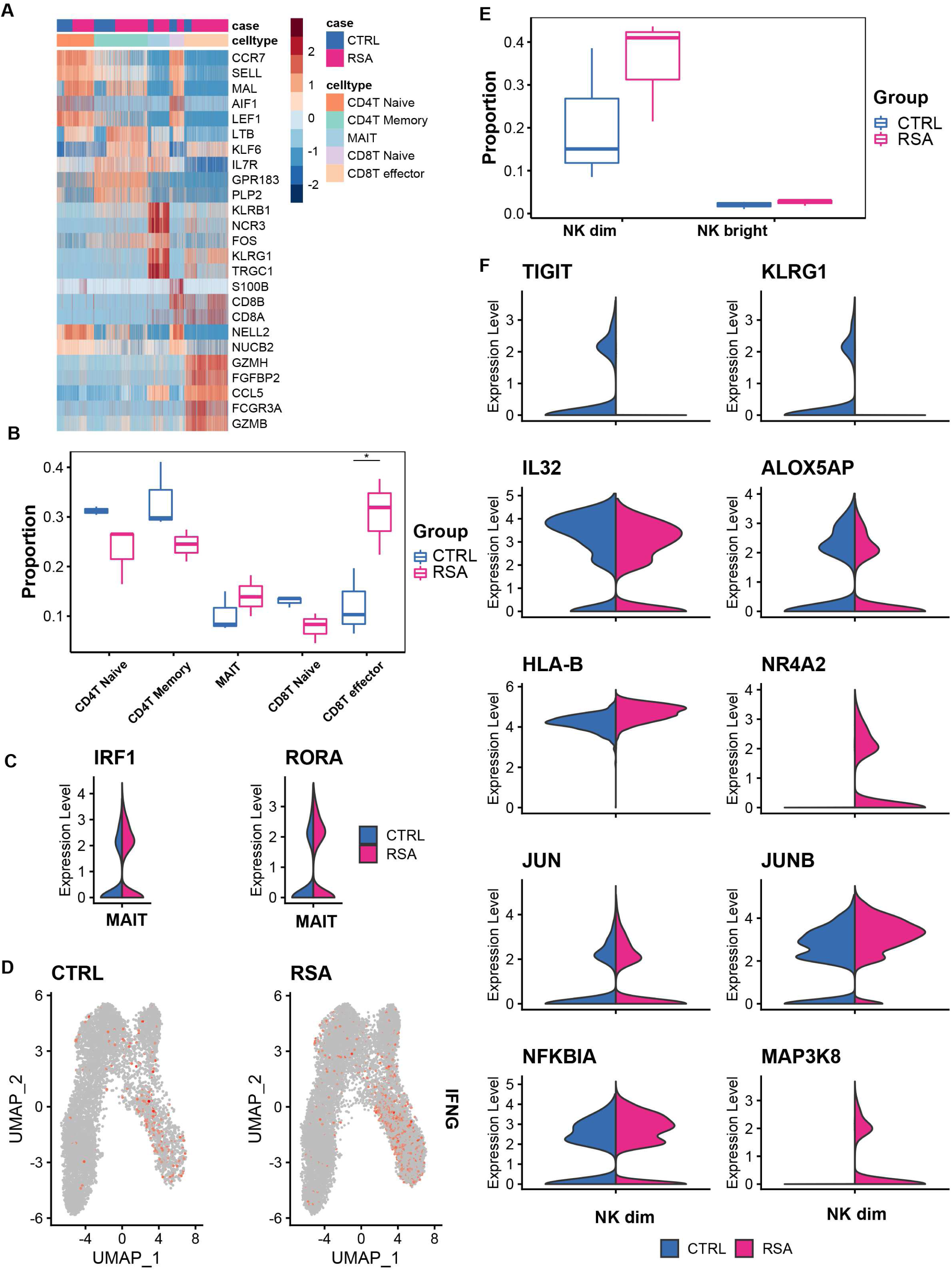
Characteristics of the peripheral blood leukocyte clusters indicate the maternal immune inflammatory status in RSA patients. (A) Heatmap of the top5 genes expressed in peripheral blood T cell subsets. x-axis represents T cell clusters; y-axis represents genes. (B) Box plots depicting cell composition of peripheral blood T cells in healthy control and RSA patients. (C) Violin plots of selected markers identified by differential expression analysis comparing the MAIT subset between RSA and CTRL patients. (D) UMAP visualization of the log-transformed, normalized IFNG expression showing the relatively stronger cytotoxicity of CD8 T effector in RSA pregnancy. high expression is showed in red, low expression is illustrated in grey. (E) Box plots depicting cell number ratio of peripheral NK cell subsets in peripheral blood of normal and RSA pregnancies. (F) Violin plots of indicated genes identified by differential expression analysis of NK dim subset between RSA and CTRL patients. Differential expression analyses in Fig. 2 were performed using the non-parametric two-sided Wilcoxon rank sum test in Seurat.

The other significant change in RSA peripheral leukocytes was the increased proportion of CD56dimCD16+ NK dim cells (Fig. 2E). Furthermore, in these cells, evidently decreased expression of immunosuppressive genes, such as *TIGIT, KLRG1, IL32* and *ALOX5AP*, while increase in pro-inflammation genes including *HLA-B, NR4A2, JUN, JUNB, NFKBIA* and *MAP3K8* were observed in RSA cases (Fig. 2F). Take together, the properties of the peripheral leukocytes, including aforementioned cell composition shift and specific gene expression pattern in certain cell cluster, strongly indicate an overall enhanced systematic pro-inflammatory immune feature of RSA patients.

### Cellular atlas of the decidual leukocytes in RSA pregnancy

Increasing evidence has demonstrated compromised immune cell differentiation or function in RSA decidua. However, a global cellular change remains largely undefined. Our analysis of cell subset proportion in decidua clearly revealed an increased portion of dCD8 T, dNKd, and dNKe populations, and lowered ratio of dNKa, dNKc, and dM cells in RSA decidua (Fig. 3A). To systematically investigate the cell-cell communications among these cell subsets, we analyzed the expression levels of immune cell revenant ligand-receptor interacting pairs within cell types (Fig. 3B and C). Significantly increased expression of *IFNG* and *TNF* was found in dCD8 T, ILC3 and dM in RSA decidua, which may account for the accumulation of dNKe or dCD4T cells that exhibited remarkably higher expression of the corresponding receptors for these two cytokines. The receptor for *CXCL16*, i.e. *CXCR6*, was predominantly expressed in dCD4T and dCD8T. Therefore, the accumulation of T cell subsets in RSA decidua may attribute to the enhanced *CXCL16* in dM. In addition, the expression of *CCL3, CCL4*, and *CCL5* in different immune cell subsets also reflected the activation of the immune micro-environment at the maternal-fetal interface and the vigorous recruitment of macrophages (Du et al., 2014). Such molecular interactions among cell populations via specific ligand-receptor complexes indicated a potential mechanism of immune activation that caused compromised immune environment in RSA decidua.

**Fig. 3.**
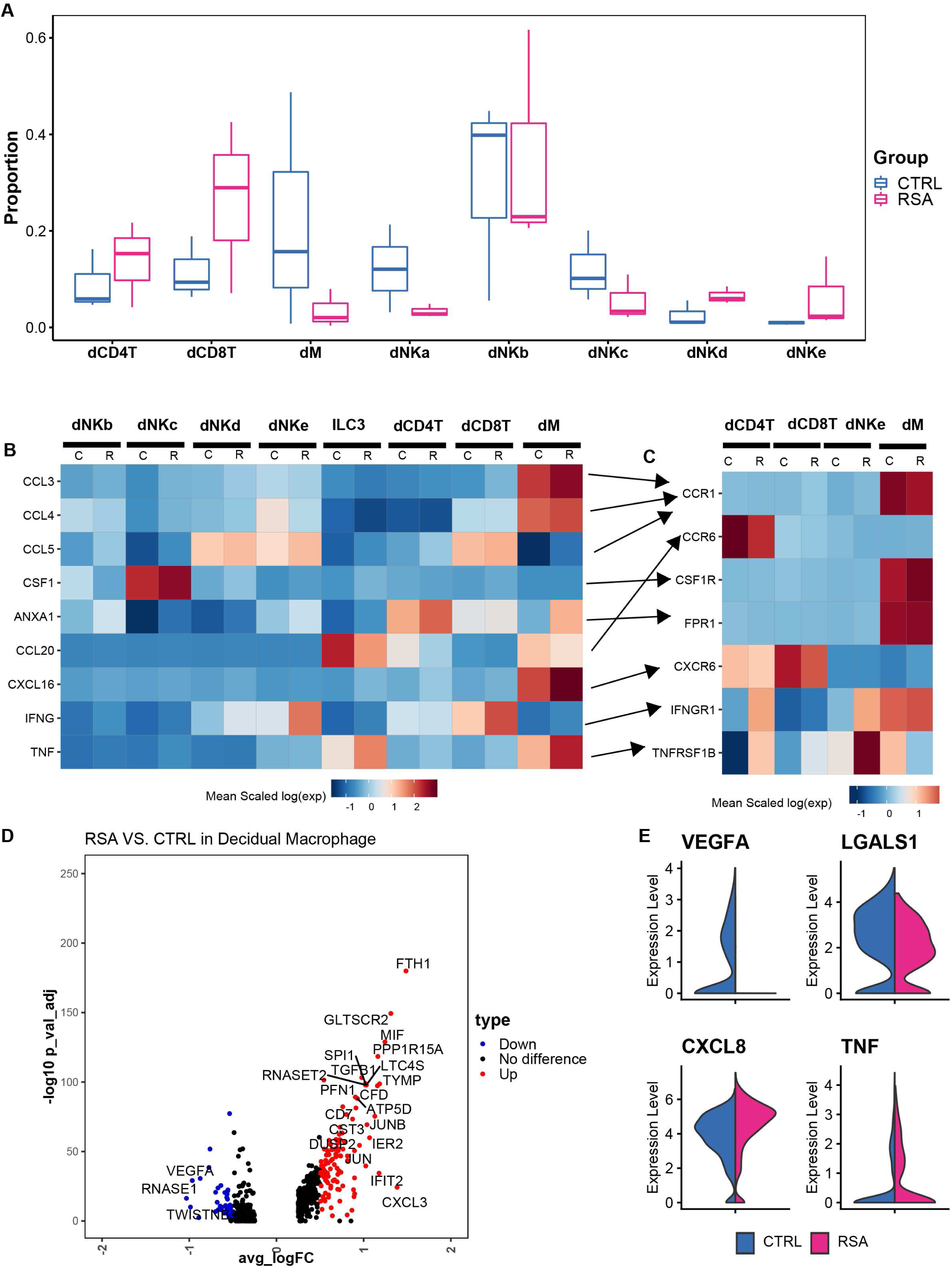
Characteristics of decidual leukocyte subpopulations demonstrated an immune activation status at the maternal-fetal interface of RSA pregnancy. (A) Cell proportion of T, dM and dNK cell subsets in decidual tissues of normal and RSA pregnancies. (B) and (C) Analysis of the ligand-receptor interactions among the leukocyte subpopulations between RSA and normal pregnancies in decidua. C: CTRL, R: RSA. (D) Volcano plot showing globally differential genes in decidual macrophages between normal and RSA pregnancies. (E) Expression of anti-inflammation and inflammation -associated cytokines in decidual macrophage subpopulation in normal and RSA pregnancies. Differential expression analyses in D were performed using the non-parametric two-sided Wilcoxon rank sum test in Seurat. Panels

### Differential gene expression profiles in decidual macrophage of RSA pregnancy

Alteration in global gene expression profiles in the RSA leukocyte subset was typically observed in dM. As shown in the volcano plot, using fold-change (FC) >1 and p-values <0.05 as cutoffs, 96 upregulated genes and 27 downregulated genes were identified in RSA dM relative to its normal counterpart (Fig. 3D).

Decidual macrophages mediate multifunctional immunoregulation, such as antigen presentation, phagocytosis, recruitment of other immune cells (Varol et al., 2015)(Gentek et al., 2014). The increased expression of *CXCL8, TNF*, and *IFIT2* genes in RSA dM indicated the impaired immune protection. Moreover, the suppressed level of angiogenic factor *VEGFA* and the immunosuppressive molecule *LAGLAS1*, while enhanced expression of pro-inflammatory factors *CXCL8* and *TNF* in RSA dM cell were observed (Fig. 3E). These data indicated a preferentially pro-inflammatory property of dM in RSA patients.

### Molecular aspects and developmental trajectories of human dNK subtypes

Consistent with the findings in previous literatures (Bulmer et al., 1991; King et al., 1991), dNK cells accounted for more than 70% of total decidual leukocytes in our dataset. The characteristics of the five dNK subpopulations were further illustrated by the top 20 marker genes (Fig. 4A) and KEGG analysis (Fig. 4B). As shown, dNKa cells highly expressed *MCM5, STMN1*, and *PCNA*, indicating their manifest proliferative capacity. The defining markers for dNKb cells included *XCL1, XCL2*, and *ZNF683*. dNKc cells strongly expressed *ENTPD1* (CD39), *CSF1* and *PKM*, and showed active oxidative phosphorylation and glycolysis activity. dNKd cells were defined by *ITGAE* (CD103), *CD160*, and *KLRB1*, and exhibited certain cytotoxic and immune-activated property. *FCGR3A*/CD16, *CXCR4, PLAC8* were enriched in dNKe cells, which indicates an active cytotoxicity status of these cells.

**Fig. 4.**
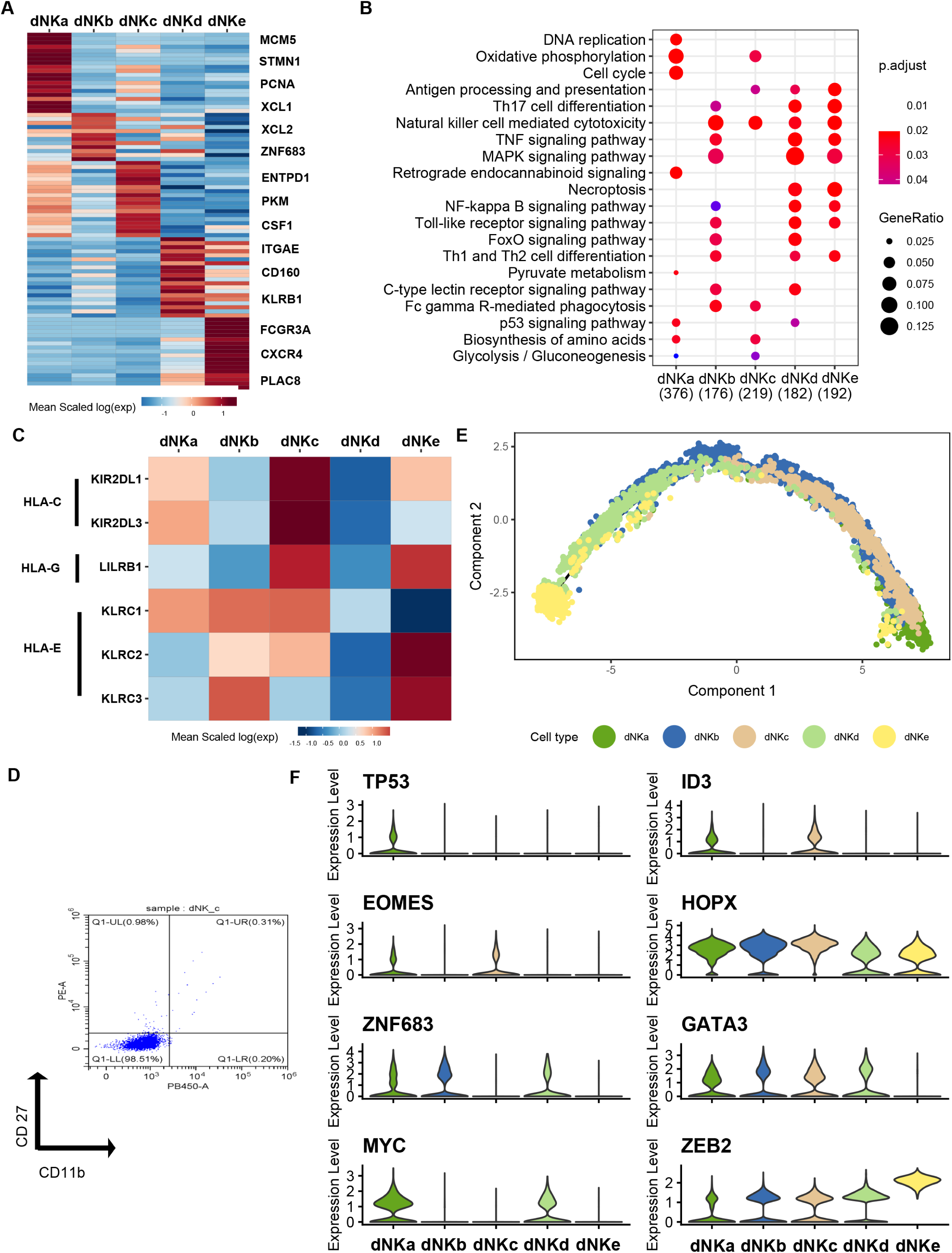
Molecular aspects and developmental trajectories of human dNK subtypes. (A) Heatmap showing relative expression (z-score) of 20 marker genes defining the subpopulations of dNK cells. (B) Functional KEGG enrichment analysis illustrating the functional signature of the dNK subpopulations. (C) Expression of KIR molecules mediating in the indicated dNK subpopulations. (D) Flow cytometry analysis showing the dNKc cells do not express CD27 and CD11b, further demonstrating their low-differentiation status. (E) Monocle2 analysis showing the developmental trajectories of dNK cells. (F) The expression of classical transcription factors in dNK subpopulations indicating their different developmental status.

It has been well known that dNK cells possess various receptors for recognition of HLA-C, HLA-G and HLA-E molecules in placental trophoblast cells, which primarily mediates dNK-trophoblast communications. We found that dNKc cells expressed much high level of *KIR2DL3* and *KIR2DL1* which are inhibitory KIR receptor for HLA-C, and *LILRB1*, the high-affinity receptor for HLA-G. dNKc and dNKb cells also moderately expressed *KLRC2*/NKG2C, *KLRC3*/NKG2E and *KLRC1*/NKG2A, which are receptors for HLA-E. dNKd cells possessed much lower expression of all these receptors. Moderate levels of *LILRB1, KLRC2*/NKG2C and *KLRC3*/NKG2E were observed in dNKe cells (Fig. 4C).

To determine whether the dNK subtypes exhibit cell type dependent localization pattern at the feto-maternal interface, we performed immunofluorescent staining for CD56, *ENTPD1*(CD39), *ITGAE*(CD103), CD16 and PLAC8 on serial sections of decidual tissues. Immunohistochemistry for cytokeratin 7 (CK7) was used to locate trophoblast and uterine gland epithelium in the sections. The results showed that CD39 positive dNKc cells were more abundant in decidual compacta where trophoblasts infiltrate, while CD103-labelled dNKd cells mainly distributed in distal compacta through spongiosa, but less in proximal compacta (Fig. 5A, B and C). A very small number of CD16-expressing dNKe cells could be found in decidual compacta, usually near uterine spiral artery (Fig. 5D). These results predict the potential of dNKc cells in the recognition and protection of trophoblasts, and dNKd cells in maintaining certain cytotoxic and immune-active environment in decidua.

**Fig. 5.**
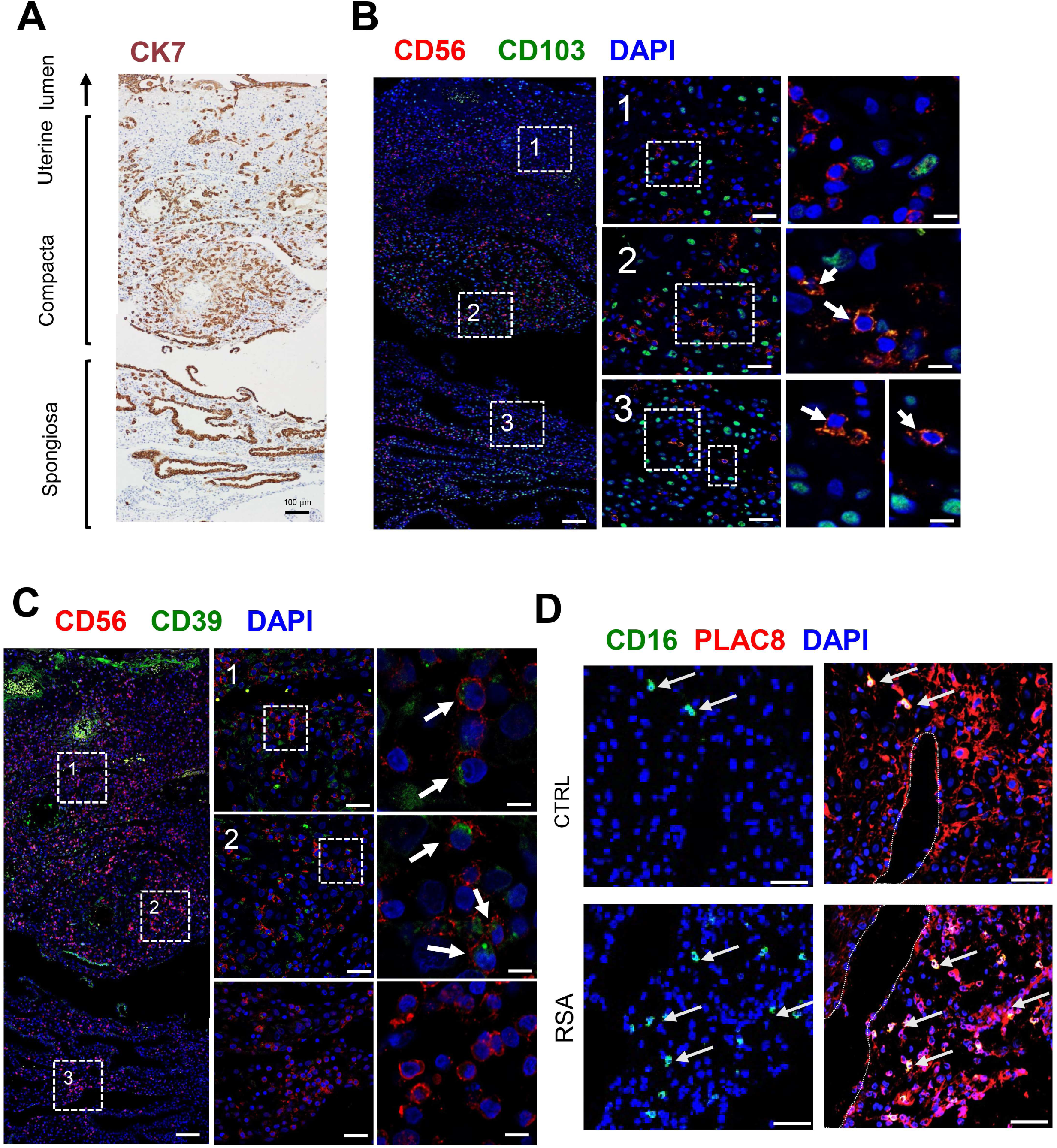
Immunofluorescence to show the spatial distribution of dNK cells at the feto-maternal interface. (A) Immunohistochemistry for cytokeratin 7 (CK7) was used to locate trophoblast and uterine gland epithelium in the sections. (B) and (C) Immunofluorescence staining show the spatial distribution of dNKc (CD39+) and dNKd (CD103+) cells at the feto-maternal interface at early pregnancy. Scale bar in panel B and C from left to right are 100, 25, 10 μm. (D) The spatial distribution of dNKe in decidual tissues from normal and RSA pregnancies by Immunofluorescence staining. Scale bar of panel D, 25μm. The arrow marks the target cell we zoomed in Fig. 5 B, C and D. The dotted line in panel D indicates the position of blood vessels

Interestingly, apart from dNKe, the other four dNK subtypes co-expressed the tissue-resident markers CD49a/*ITGA1*, indicating their uterine resident feature. By far the developmental route of uterine resident dNK cells remains unclear. Our result of flow cytometry showed dNKc cells as CD27-CD11b-(Fig. 4D), which indicates their immature developmental status according to the previous report (Fu et al., 2011). Therefore, we further analyzed the developmental trajectory of dNK cells by carrying out pseudotime analysis on our scRNA-seq data. The top 100 genes in each dNK subtype were clustered (Fig. S4), and Monocle2 was performed to infer their development trajectory (Fig. 4E). As shown, dNKa appeared at the very initiation of the trajectory, and dNKc emerge at the early developmental stage with much higher expression of dNK-classical transcription factors *EOMES* and *ID3*. It tended to differentiate towards dNKb and finally dNKd (Fig. 4F). Distance between dNKd and dNKe indicated the probably alternative origin of CD49a-CD16+ dNKe cells.

### Alterations of dNK subsets in decidua of RSA pregnancy

To verify the feature of dNK subset proportion in RSA decidua identified by scRNA-seq in Fig 3A, we analyzed dNK cell population by flow cytometry in a larger sample cohort of normal and RSA decidual tissues. We sorted cells by using CD56 (expressed by all dNK cells) and CD3 (negative sorting marker), combined with markers for dNKc (CD39), dNKd (CD103), and dNKe (CD16) that were identified from our scRNA-seq data (Fig. 6A). As indeed, the cell portion of dNKc was uniformly and remarkably reduced by around 50%, whereas dNKd proportion increased by 1-fold in RSA decidua, compared to the normal pregnancy controls (Fig. 6B). The ratio of dNKe cells was rather small in normal decidua, accounting for less than 4% of total dNK. However, in approximately half of the enrolled RSA cases, dNKe cell population increased to about 12% of total NK cells (Fig. 6B). Consistently, immunofluorescent staining for CD16 in RSA decidua showed evidently more dNKe cells in decidua compacta niche close to blood vessels (Fig. 5D).

**Fig. 6.**
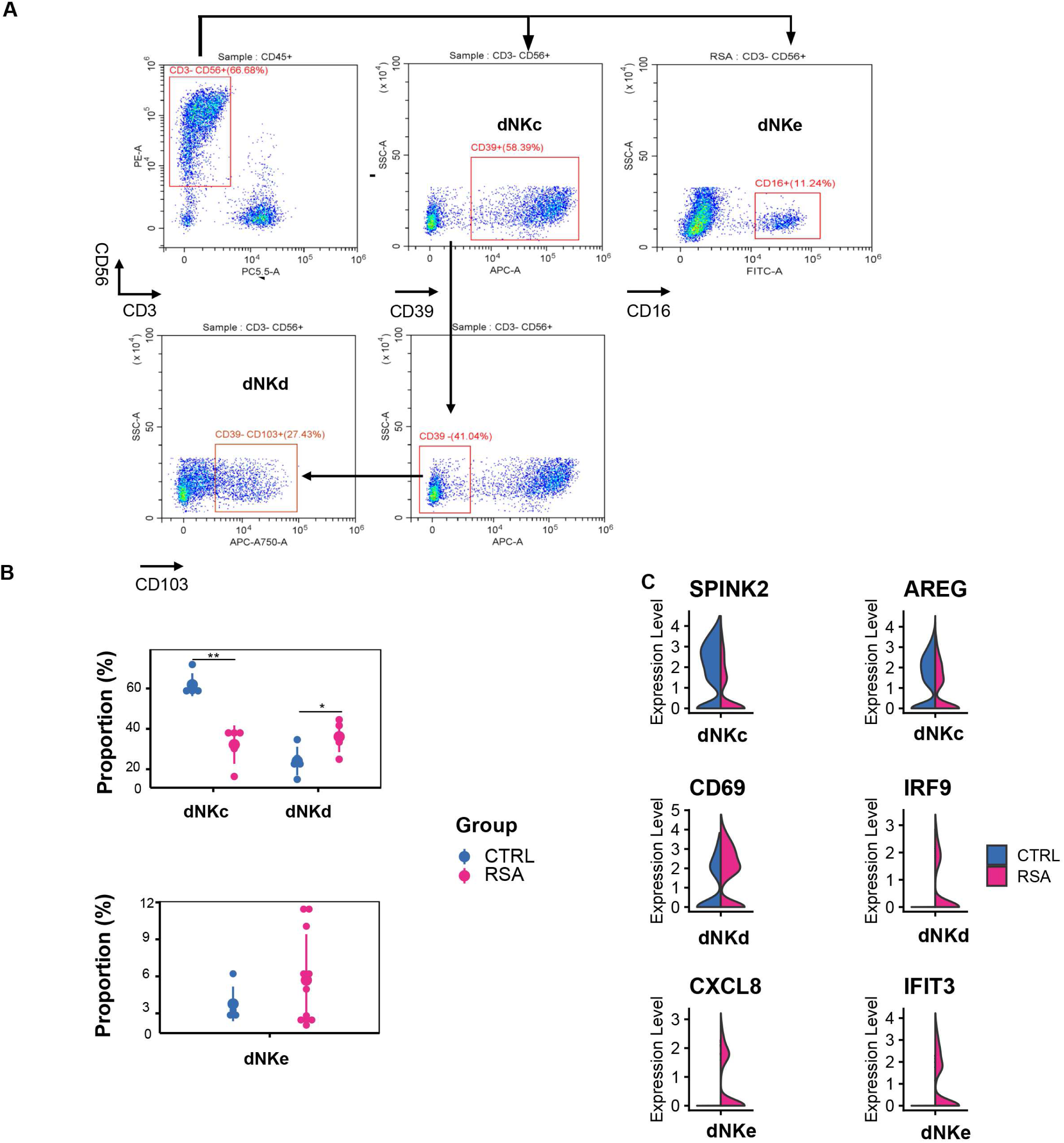
Alterations of dNK subsets in decidua of RSA pregnancy. (A) Flow cytometry to sort dNKc (CD39+), dNKd (CD103+) and dNKe(CD16+) cells in human decidual tissues. (B) Cell population of dNKc and dNKd in decidua tissues from normal pregnancy (n=5) and RSA (n=5). Lower panel, Statistical results showing cell proportions of dNKe in decidua tissues from normal pregnancy (n=4) and RSA (n=11). (C) Violin plots revealing differential expression of indicated genes in dNKc, dNKd and dNKe subsets from RSA and CTRL patients. Significance was analyzed using unpaired t-tests assuming no consistent s.d..

Comparison of the differential gene expression in dNK subsets between RSA and normal pregnancy demonstrated a significant increased expression of inflammation-related genes such as *CD69* and *IRF9* in dNKd cells, *INFG, CXCL8, TNFRSF8*, and *IFIT3* in dNKe cells, as well as downregulation of anti-inflammation genes including *SPINK2* and *AREG* in dNKc cells from RSA patients (Fig. 6C).

## DISCUSSION

Cellular and molecular mechanisms accounting for maternal immune tolerance to protect growth of semi-allogenic fetus in placental mammals has been a fascinating scientific question. Increasing evidence demonstrate the significance of the complicated and precisely controlled cell-cell communications within the feto-maternal interface in concert with the well-coordinated systematic immune adaptations along gestation (Ander et al., 2019; Arck and Hecher, 2013; Erlebacher, 2013; Hanna et al., 2006; Nancy et al., 2012). Recently, a molecular and cellular map of normal feto-maternal interface at human early gestation strikingly deepened our understanding of the cell subtypes and their predicted interactions in normal homeostatic decidua (Vento-Tormo et al., 2018). However, the corresponding landscape and dynamics of immune cell at single cell level in failed pregnancy, such as RSA, is lacking to date. To our knowledge, our study is the first comprehensive single-cell transcriptomics atlas of the decidual and peripheral immune cells in human recurrent miscarriage at early gestation. Our findings integrate complex information of immune cell, including cell composition, functional status, developmental trajectory, and predicted cellular interactions among immune cells. Our work illustrates an integral framework of the compromised immune environment that is harmful to the intra-uterine fetus in RSA pregnancy.

The immuno-mechanism for RSA, especially those without fetal chromosomal or congenital abnormalities or other known pathological causes, has been largely debatable. Multiple studies demonstrated abnormal expressions or functions of various cytokines, or dysregulated proportions or properties of certain immune cells in patients with recurrent miscarriage. However, studies solely focusing on certain limited cell types may result in misleading points or contradictive results. To date the strategies of early diagnosis or intervention of RSA are lacking, which makes it hard to reduce the threat to the patients (Seshadri and Sunkara, 2014). Here by taking advantage of single-cell sequencing technology, we construct an overall pathological change in the properties of immune cell subsets in peripheral blood, which may pave the path for the pre-symptom diagnosis of the disordered pregnancy. The significantly enhanced differentiation of naive T cells to CD8T effector cells, and preferential increasing in number of NK dim cells, together with the upregulated inflammation-related genes, demonstrate a systematic pro-inflammatory status in RSA patients. These results are consistent with previous reports (Ebina et al., 2017; Kuon et al., 2017). An interesting finding is the increased frequency of MAIT cells in RSA peripheral blood. As a non-conventional T cell subset, MAIT cells are mainly activated by exposure to microbes, while they can also be turned on by inflammatory stimuli in the absence of TCR-mediated antigen recognition. There is considerable evidence to suggest their involvement in a broad range of infectious and non-infectious diseases (Godfrey et al., 2019). Studies in liver disorders suggest that the MAIT cells may play a protective role against bacterial infections in a normal liver, but might be detrimental, with over-inflammation, in liver diseases (Bertrand and Lehuen, 2019). Notably, MAIT cells express several cytokine receptors, including IL-1R, IL-7R, IL-12R, IL-15R, IL-18R and IL-23R, and response to these multiple cytokines stimulation (Godfrey et al., 2019). Our sequencing data showed the evidently enhanced expression of *IRF1* and *RORA* in MAIT from RSA periphery, indicating their highly immune-activated property. By far evidence of MAIT cell in pregnant condition has been very limited, and our study for the first time indicate the association of MAIT activation with the maternal inflammation condition that damage fetal survival. Whether the change of MAIT activation is the cause or result of miscarriage remains to be established.

At the maternal-fetal interface, decidual NK cells assemble the largest population of decidual leukocytes at early pregnancy. Unlike their peripheral counterparts, dNK cells are less cytotoxic but actively secret a vast array of factors and cytokines. They have been found to play powerful roles including facilitating the remodeling of uterine spiral arteries, promoting trophoblast invasion and fetal growth, regulating T cell differentiation, increasing the availability of maternal blood at the implantation site, and so on (Ander et al., 2019). In unexplained infertile patients, substantially fewer uterine NK cells were observed when compared to fertile controls (Klentzeris et al., 1994). However, discrepant results have been reported regarding the change of dNK cell number in RSA patients (King et al., 2010; Seshadri and Sunkara, 2014). Our study clearly demonstrates the abnormal properties of dNK subsets in RSA decidua, including cell composition and gene expression, which suggests impaired regulation of dNK subset development in the patients.

The characteristics of dNKc cells are similar to the highly active dNK1 subset in Vento-Tormo’s study. They are numerically dominant dNK cells and represent the typical dNK functions to support embryo growth. The CD11b-CD27-property of dNKc manifests their immature characteristic and the potential to differentiate (Fu et al., 2011). Specific expression of PBX1 was found in CD27-CD11b-dNK cells, and a recent study demonstrated the association of decreased PBX1 expression or PBX1^G21S^ mutation with unexplained RSA (Zhou et al., 2020), further suggesting the crucial role of this dNK subset in pregnancy maintenance. Interestingly, a recently identified pregnancy trained dNK (PTdNK) share similar features with dNKc or dNK1 (Gamliel et al., 2018b), and this subset has been speculated to be enriched and boost decidua receptivity in subsequent pregnancy, therefore may be responsible for the “memory” of reproductive outcomes in next pregnancies. Thus, the strikingly lowered frequency of this subset in RSA patients can partly explain why failed pregnancies repeatedly occur in the patients. In addition to the decreased proportion, the altered gene expression in dNKc of RSA decidua further indicates their diminished immune-protective capability.

The development trajectory of dNK subsets reveals dNKd cells are relatively mature and immune-activated. They are likely to maintain certain cytotoxic and immune-active environment in decidua, which may contribute to the appropriate extent of trophoblast invasion. However, in RSA decidua their portion raises to more than one fold of the normal counterpart. Besides, they exert enhanced production of pro-inflammatory cytokines, which is likely causing the decidual environment harmful to the fetal cells and other maternal immune cells such as T cells and macrophages (Fridman et al., 2012; Fu et al., 2013; Ma et al., 2017; Sotnikova et al., 2014; Yang et al., 2019).

Another notable dNK subset in our study is the newly-identified CD56+CD16+ dNKe, which is largely enriched in RSA decidua. Being different from all the other four subsets, dNKe cells do not exert expression of the tissue-resident marker CD49a. Consistently, the developmental trajectory show separation of dNKe from the maturation route of other dNK subsets. It is therefore likely that this subset may origin from other sources. Currently there are three hypotheses for the origin of dNK cells, i.e., 1) recruitment of peripheral blood NK cells to decidua(Carlino et al., 2008), 2) maturation of uterine-resident NK cells in response to IL15 or progesterone(Manaster et al., 2008), 3) direct differentiation from hematopoietic precursors in the decidua upon stimulation of specific decidual factors(Vacca et al., 2011). Considering the common CD49a-CD16+ feature of dNKe and peripheral NK dim, we further compared the gene expression pattern between dNKe and NK dim. As shown in the volcano plot, using fold-change (FC) >1 and p-values <0.05 as cutoffs, 187 upregulated genes and 73 downregulated genes were identified in dNKe relative to NK dim. However, there are 699 co-expressed genes between dNKe and NK dim, which present their common feature in exercising common cellular functions (Fig. S5). Thus, it is likely that peripheral NK dim cells are recruited to decidua and further educated towards dNKe by some decidual factors. Many studies have demonstrated the crucial role of IFN-gama, CCL3/MIP-1, CXCL10/IP-10, CXCL12/SDF-1 in enrolling pNK to decidua (Hanna et al., 2003; Wallace et al., 2013; Wu et al., 2005). What’s more, an in vitro study reveals that human peripheral CD16+ NK cells can be converted to a dNK-like phenotype upon the stimulation of hypoxia, TGF-beta, and the demethylating agent Aza (Zhang et al., 2016). Here our data reveals the evident increase in *INFG* in dCD8T, *CXCL16, CCL3* and *TGFB* in dM of RSA decidua, which may be responsible for enrolling and educating excessive pNK cells to decidua compartment. The much higher expression of several pro-inflammatory factors in RSA dNKe cells also indicate their enhanced cytotoxicity in the patients. An interesting previous study demonstrated the extravagant enrichment of CD16+ NK cells in endometrium of RSA patients during their pre-pregnant period (Lachapelle et al., 1996). Therefore, we propose that the patients with more abundant dNKe cells may suffer a greater chance of pregnancy failure in their subsequent pregnancies.

Decidual macrophages account for around 20% of leukocytes at the feto-maternal interface, and they have many diverse functions during pregnancy. Here a relatively large amount of differentially expressed genes was identified in RSA decidual macrophages, which potentially suggest some functional abnormalities. For instance, the obviously repressed *VEGFA* in RSA dM indicates the link to impaired remodeling of the spiral arteries and angiogenesis. The upregulated genes include immunoinflammatory factors and the relevant signaling molecules, such as *CXCL8, TNF, IFIT2, JUN, JUNB*, etc., predicting the diminishment in the anti-inflammatory capacity. Furthermore, the increased expression of IL-8 in dM and enhanced *IFNG* expression in dNKe of RSA patients are likely in tight correlation, because it has been demonstrated that IL-8 from dM enhances the production of IFN-gama in dNK cells (Baratin et al., 2005; Dalbeth et al., 2004). In addition, our brief repository of ligand-receptor complexes indicated the potential of dM in recruiting dCD8T or dNKe through *CXCL16*-*CXCR6* or *TNF*-*TNFRSF1B* interactions. Although decidual macrophages are believed to exist predominantly in a regulatory/homeostatic M2-like (dM2) phenotype, while less in pro-inflammatory M1 (dM1) phenotype during pregnancy(Erlebacher, 2013), we did not find an alteration in the proportion of dM1 and dM2 in RSA patients (Fig. S6). This is probably due to the relatively small amount of captured dM cells for sequencing.

It has to be noticed that the subset analysis on some low-abundant cell subpopulations, such as decidual T cells, DC cells and macrophages, remains insufficient. Therefore, it warrants further investigation on the subset characterization of these cells by specifically improved enrichment strategy, which will provide a more comprehensive understanding of the complicated cell-cell communication network at the feto-maternal interface. In addition, the analysis of endometrial cell atlas in the pre-pregnant stage of RSA patients will be helpful to find prospective intervention targets and improve their pregnancy success.

In general, our study comprehensively illustrates the compromised immune response in periphery and feto-maternal interface of RSA patients. The findings generate a data-driven hypothesis about immune-related pathogenesis for recurrent miscarriage, and provide new insight into the strategies for diagnosis and intervention of the diseases.

## MATERIALS AND METHODS

### Sample Collection and Ethic permission

Clinical samples of anti-coagulant peripheral blood and decidual tissues from normal pregnancy (N=10) or recurrent spontaneous abortions (RSA) (N=14) at gestational week 6-8 were obtained upon therapeutic termination of pregnancy at Peking University Third Hospital. The decidual tissues were immersed in iced RPMI-1640 medium and the blood samples were kept on ice. All samples were subjected to cell isolation or fixation within 1 hour following the surgery. The collection of human samples was permitted by the Local Ethical Committees in Peking University Third Hospital.

RSA was defined according to the criteria of Practice Committee of the American Society for Reproductive Medicine. In brief, these patients had history of two or more failed pregnancies with unknown cause(Committee and Society, 2013). Women who manifest endocrine disorder, fetal chromosomal or congenital abnormalities, uterine anatomical disorders, renal disease, or pregnancies conceived by fertility treatment were excluded from this study. All the enrolled patients had arrested fetal development for less than one week before the termination of pregnancy.

### Cell isolation and purification

Freshly collected human decidual tissues were trimmed into 1 mm3 piece by GentelMACS Dissociator (Miltenyi Biotec, 130-093-235, Germany) and digested twice for 30 minutes each at 37°C with 1.0 mg/ml type IV collagenase (Gibco, 9001121, Grand Island, NY, USA) and 10U/ml type I DNase (Sigma, DN25, St Louis, MO, USA). The cell suspensions were filtered through 60 mesh and 200 mesh sieves, and were collected by centrifuging at 1000rpm for 10 min. The resuspended cells were subjected to lymphocyte enrichment using Ficoll-Paque Plus (17-1440-02, GE Healthcare). The decidual leukocytes were further purified by fluorescent activated cell sorting with 7-AAD (420404, Bioledgend) and FITC-labeled anti-CD45 antibody (11-0459-42, eBioscience).

Freshly collected peripheral blood samples were diluted to 1:1 with PBS. Which Peripheral blood after dilution were subjected to lymphocyte enrichment using Ficoll-Paque Plus (17-1440-02, GE Healthcare). The peripheral blood leukocytes were further purified by fluorescent activated cell sorting with 7-AAD (420404, Bioledgend) and FITC-labeled anti-CD45 antibody (11-0459-42, eBioscience).

The freshly purified decidual or peripheral leukocytes were immediately subjected to single cell RNA sequencing as described below.

### Generation of single-cell library and transcriptome sequencing

The purified leukocytes from three pairs of normal and RSA cases were separately loaded on Chromium Single Cell Controller (10x Genomics) using the Chromium Single-cell 3’ kit v2 to capture 5000 to 8000 cells per sample.

Libraries were sequenced on an Illumina NovaSeq 6000 (Illumina, San Diego) with a read length of 26bp for read 1 (cell barcode and UMI), 8 bp i7 index read (sample barcode), and 98 bp for read 2 (actual RNA read). Reads were first sequenced in the rapid run mode, allowing for fine-tuning sample ratios in the following high-output run. Combining the data from both flow cells yielded approximately > 40,000 reads per cell.

### Single-cell RNA-seq data processing and analysis

The raw sequencing reads were processed using Cell Ranger (version 2.0.1, 10x Genomics) (Zheng et al., 2017) The reference index was built using the GRCh38 human reference genome assembly. Cells with fewer than 600 detected genes or with the total mitochondrial gene expression exceeded 5% were removed. Then we convert the obtained matrix into a Seurat object for downstream analysis. All Seurat objects for individual samples were merged into one combined object. According to the integration method reciprocal PCA provided by Seurat, we first performed standard normalization and variable feature selection on each individual sample. Next, we selected features for downstream integration, and ran PCA on each individual sample in the combined object. Since our data include RSA patients and normal pregnant women, we chose one of RSA and normal pregnant women data as the reference, and then used the function FindIntegrationAnchors provided by Seurat to integrate the samples of 6 individuals. Downstream analyses, including normalization, shared nearest neighbor graph-based clustering, differential expression analysis, and visualization, were performed using the standard workflow provided by Seurat (version 3.0.3). Developmental trajectories were inferred with the Monocle2 (version 2.12.0),. We used the function FindVariableGenes of Seurat to find the variant genes of dNK cells, and then performed trajectory analysis based on the standard workflow provided by Monocle2. We performed differential gene expression analysis of all decidual cell types using Wilcoxon rank sum test inbuilt in Seurat package. differentially expressed genes for dNK clusters were KEGG analysis using R package clusterProfiler(version 3.12.0)(Yu, 2015).

### Immunostaining analysis

Human decidual tissues were briefly fixed in 4% PFA and embedded in O.C.T. compound (Sakura Finetek, Torrance, CA, USA). The frozen sections at 10 um were further fixed in 4% PFA and treated with 0.1% triton, and subjected to the incubation with specific antibodies against NCAM1/CD56 (Abcam, ab75813, 1:300), CK7 (Abcam, ab181598, 1:10000), CD39 (proteintech, 14211-1-AP, 1:200), or CD103 (350227, Biolegend, 1;200). Binding of the antibody was visualized using FITC-conjugated or TRITC-conjugated secondary antibody (ZSGB-BIO, ZF-0311, ZF-0313, 1:100), and cell nuclei were stained with 4, 6-diamidino-2-phenylindole (DAPI; Sigma, 28718-90-3, St Louis, MO, USA). The results were recorded using Zeiss LSM780 confocal system (Zeiss, Oberkochen, BW, Germany) and processed with ZEN 2012 software (Zeiss). Immunohistochemical staining for Cytokeratin (CK) in decidua was performed by using antibody against CK7 (ab181598, Abcam,1:10000) and HRP-conjugated second antibody (ZSGB-BIO, PV-6001) followed by recovery of substrate DAB (ZSGB-BIO, ZLI-9019). The imagines were recorded on a light microscope with CCD (DP72, Olympus, Japan).

### Flow cytometry Assay

Flow cytometry assay for dNK cells was carried out in CytoFLEX (Beckman Coulter, lnc.) using the following antibodies: PE-labeled anti-CD56(362508, Bioledgend), Percp-cy5.5-labeled anti-CD3(300328, Bioledgend), APC-labeled ani-CD39(328209, Bioledgend), APC-cy7-labeled anti-CD103(350227, Bioledgend), FITC-labeled anti-CD16(302206, Bioledgend), PE-labeled anti-CD27(356405, Bioledgend), Pacific blue-labeled anti CD11b(301316, Bioledgend) according to the manufacturer’s instructions. Data was analyzed using CytExpert (Beckman Coulter, lnc.).

### Statistical analysis

Comparison of cell proportion between normal and RSA pregnancies were statistically analyzed with GraphPad Prism version 7.00 (GraphPad Software, San Diego, CA, USA). Data were shown as Mean±S.E.M. and comparison was carried out by unpaired t-test. The P values of less than 0.05 were considered statistically significant. All the differential expression analyses were performed using the non-parametric two-sided Wilcoxon rank sum test in Seurat.

### Data and code availability

The raw sequence data reported in this paper have been deposited in the Genome Sequence Archive(Wang et al., 2017) in National Genomics Data Center(Zhang et al., 2020), Beijing Institute of Genomics (China National Center for Bioinformation), Chinese Academy of Sciences, under accession number HRA000237 that are publicly accessible at http://bigd.big.ac.cn/gsa-human.

## ACKNOWLEDGEMENTS

We are grateful to Prof. Bin Cao at Xiamen University and Mr. Zhenghui Zhao at IOZ for their critical comments to this study. The technical support from Shiwen Li, Xia Yang and Qing Meng in the experiments of Confocal analysis and FACS was appreciated. We acknowledge all the enrolled patients for their contribution to the study.

This study was supported by grants from the National Key Research and Development Program of China (2018YFC10041002, 2017YFC1001404, 2016YFC1000401, 2016YFC1000200), and the National Natural Science Foundation (81730040, 81490740).

## AUTHOR CONTRIBUTIONS

F.W. performed cell isolation, flow cytometry and scRNA sequencing data analysis, and drafted manuscript. W.J. carried out immunostaining and flow cytometry, and participated in data analysis. M.F. was in charge of patient enrollment, sample collection and clinical data checking. Y.L. participated in scRNA sequencing data analysis and manuscript revision. Z.L. helped in immunostaining and flow cytometry. Y.M., X.S. and Y.L. were involved in data analysis and manuscript drafting. Y.L.W., Q.T. and R.L. designed and supervised the study, interpreted the data and revised the manuscript.

---

Fig. S1. Gating strategy and quality control for droplet single-cell RNA-seq.

(A) Flow Sorting of living leukocytes from peripheral blood and decidual tissue using CD45 antibody and 7AAD. (B) Workflow to depict the dissociation and sorting of CD45+ leukocytes from peripheral blood and decidual to generate scRNA transcriptome profile. (C)Total number of cells that met the requirement of quality control and subjected to droplet scRNA-seq. (D) Quality evaluation on sequencing data including nFeature_RNA and nCount_RNA in peripheral blood and decidua. E, Every cluster distribution of each cell subpopulation in the 6 samples.

Fig. S2. The MNN algorithm to confirm the appropriate elimination of the batch effect of the data.

(A) Data from the 6 samples were integrated with MNN and visualized using UMAP. (B) Distribution of leukocytes from peripheral blood and decidual tissues after MNN integration. (C) Mutual nearest neighbors (mnn_1 and mnn_2) of the integrated analysis of decidual and peripheral blood leukocytes from the droplet-based datasets. (D) Distribution of leukocytes from normal pregnancy and RSA after MNN integration.

Fig. S3. Identification of peripheral blood T cell subpopulations by specific marker genes.

(A) UMAP visualization of peripheral blood T cells. (B)-(E), Feature plot of Naïve T (B), MAIT T (C), CD4T memory (D) and CD8T effector (E) cells with specific marker genes.

Fig. S4. Top100 genes in dNK cells that change in a continuous manner over pseudotime

(A)Heatmap showing the top100 genes in dNK cells that change in a continuous manner over pseudotime

Fig. S5. Globally differential genes in CD16+ NK cell between dNKe from decidua and NK dim from peripheral blood

(A) Volcano plot shows that upregulated and downregulated genes between dNKe and NK dim. Differential expression analyses in dNKe were performed using the non-parametric two-sided Wilcoxon rank sum test in Seurat.

Fig. S6. Identification of decidual macrophage subpopulations

(A) UMAP showing two subpopulations of decidual macrophage. (B) UMAP showing macrophage subpopulations in normal pregnancy (grey) and RSA (red). (C) Characterization of the two macrophage subpopulations using marker genes.

## REFERENCES

Ander, S.E., Diamond, M.S., and Coyne, C.B. (2019). Immune responses at the maternal-fetal interface. Sci. Immunol.

Arck, P.C., and Hecher, K. (2013). Fetomaternal immune cross-talk and its consequences for maternal and offspring’s health. Nat. Med.

Baratin, M., Roetynck, S., Lépolard, C., Falk, C., Sawadogo, S., Uematsu, S., Akira, S., Ryffel, B., Tiraby, J.G., Alexopoulou, L., et al. (2005). Natural killer cell and macrophage cooperation in MyD88-dependent innate responses to Plasmodium falciparum. Proc. Natl. Acad. Sci. U. S. A.

Bertrand, L., and Lehuen, A. (2019). MAIT cells in metabolic diseases. Mol. Metab.

Bulmer, J.N., Morrison, L., Longfellow, M., Ritson, A., and Pace, D. (1991). Granulated lymphocytes in human endometrium: Histochemical and immunohistochemical studies. Hum. Reprod.

Butler, A., Hoffman, P., Smibert, P., Papalexi, E., and Satija, R. (2018). Integrating single-cell transcriptomic data across different conditions, technologies, and species. Nat. Biotechnol.

Carlino, C., Stabile, H., Morrone, S., Bulla, R., Soriani, A., Agostinis, C., Bossi, F., Mocci, C., Sarazani, F., Tedesco, F., et al. (2008). Recruitment of circulating NK cells through decidual tissues: A possible mechanism controlling NK cell accumulation in the uterus during early pregnancy. Blood.

Committee, P., and Society, A. (2013). Definitions of infertility and recurrent pregnancy loss: A committee opinion. Fertil. Steril. 99, 63.

Dalbeth, N., Gundle, R., Davies, R.J.O., Lee, Y.C.G., McMichael, A.J., and Callan, M.F.C. (2004). CD56 bright NK Cells Are Enriched at Inflammatory Sites and Can Engage with Monocytes in a Reciprocal Program of Activation. J. Immunol.

Deshmukh, H., and Way, S.S. (2019). Immunological Basis for Recurrent Fetal Loss and Pregnancy Complications. Annu. Rev. Pathol. Mech. Dis.

Du, M.R., Wang, S.C., and Li, D.J. (2014). The integrative roles of chemokines at the maternal-fetal interface in early pregnancy. Cell. Mol. Immunol.

Ebina, Y., Nishino, Y., Deguchi, M., Maesawa, Y., Nakashima, Y., and Yamada, H. (2017). Natural killer cell activity in women with recurrent miscarriage: Etiology and pregnancy outcome. J. Reprod. Immunol.

Erlebacher, A. (2013). Immunology of the Maternal-Fetal Interface. Annu. Rev. Immunol.

Fisher, S.J. (2015). Why is placentation abnormal in preeclampsia? Am. J. Obstet. Gynecol. 213, S115–S122.

Fridman, W.H., Pagès, F., Saut’s-Fridman, C., and Galon, J. (2012). The immune contexture in human tumours: Impact on clinical outcome. Nat. Rev. Cancer.

Fu, B., Wang, F., Sun, R., Ling, B., Tian, Z., and Wei, H. (2011). CD11b and CD27 reflect distinct population and functional specialization in human natural killer cells. Immunology.

Fu, B., Li, X., Sun, R., Tong, X., Ling, B., Tian, Z., and Wei, H. (2013). Natural killer cells promote immune tolerance by regulating inflammatory TH17 cells at the human maternal-fetal interface. Proc. Natl. Acad. Sci. U. S. A.

Gamliel, M., Goldman-Wohl, D., Isaacson, B., Gur, C., Stein, N., Yamin, R., Berger, M., Grunewald, M., Keshet, E., Rais, Y., et al. (2018a). Trained Memory of Human Uterine NK Cells Enhances Their Function in Subsequent Pregnancies. Immunity 48, 951-962.e5.

Gamliel, M., Goldman-Wohl, D., Isaacson, B., Gur, C., Stein, N., Yamin, R., Berger, M., Grunewald, M., Keshet, E., Rais, Y., et al. (2018b). Trained Memory of Human Uterine NK Cells Enhances Their Function in Subsequent Pregnancies. Immunity.

Gentek, R., Molawi, K., and Sieweke, M.H. (2014). Tissue macrophage identity and self-renewal. Immunol. Rev.

Godfrey, D.I., Koay, H.F., McCluskey, J., and Gherardin, N.A. (2019). The biology and functional importance of MAIT cells. Nat. Immunol.

Haghverdi, L., Lun, A.T.L., Morgan, M.D., and Marioni, J.C. (2018). Batch effects in single-cell RNA-sequencing data are corrected by matching mutual nearest neighbors. Nat. Biotechnol.

Hanna, J., Wald, O., Goldman-Wohl, D., Prus, D., Markel, G., Gazit, R., Katz, G., Haimov-Kochman, R., Fujii, N., Yagel, S., et al. (2003). CXCL12 expression by invasive trophoblasts induces the specific migration of CD16-human natural killer cells. Blood.

Hanna, J., Goldman-Wohl, D., Hamani, Y., Avraham, I., Greenfield, C., Natanson-Yaron, S., Prus, D., Cohen-Daniel, L., Arnon, T.I., Manaster, I., et al. (2006). Decidual NK cells regulate key developmental processes at the human fetal-maternal interface. Nat. Med.

Huhn, O., Ivarsson, M.A., Gardner, L., Hollinshead, M., Stinchcombe, J.C., Chen, P., Shreeve, N., Chazara, O., Farrell, L.E., Theorell, J., et al. (2020). Distinctive phenotypes and functions of innate lymphoid cells in human decidua during early pregnancy. Nat. Commun.

Kheshtchin, N., Gharagozloo, M., Andalib, A., Ghahiri, A., Maracy, M.R., and Rezaei, A. (2010). The expression of Th1- and Th2-related chemokine receptors in women with recurrent miscarriage: The impact of lymphocyte immunotherapy. Am. J. Reprod. Immunol.

King, A., Balendran, N., Wooding, P., and Loke, Y.W. (1991). Cd3– Leukocytes Present in the Human Uterus During Early Placentation: Phenotypic and Morphologic Characterization of the Cd56++ Population. Dev. Immunol.

King, K., Smith, S., Chapman, M., and Sacks, G. (2010). Detailed analysis of peripheral blood natural killer (NK) cells in women with recurrent miscarriage. Hum. Reprod.

Klentzeris, L.D., Bulmer, J.N., Warren, M.A., Morrison, L., Li, T.C., and Cooke, I.D. (1994). Lymphoid tissue in the endometrium of women with unexplained infertility: Morphometric and immunohistochemical aspects. Hum. Reprod.

Kuon, R.J., Vomstein, K., Weber, M., Müller, F., Seitz, C., Wallwiener, S., Strowitzki, T., Schleussner, E., Markert, U.R., Daniel, V., et al. (2017). The “killer cell story” in recurrent miscarriage: Association between activated peripheral lymphocytes and uterine natural killer cells. J. Reprod. Immunol.

Lachapelle, M.H., Miron, P., Hemmings, R., and Roy, D.C. (1996). Endometrial T, B, and NK cells in patients with recurrent spontaneous abortion. Altered profile and pregnancy outcome. J. Immunol.

Ma, L., Li, G., Cao, G., Zhu, Y., Du, M.R., Zhao, Y., Wang, H., Liu, Y., Yang, Y., Li, Y.X., et al. (2017). DNK cells facilitate the interaction between trophoblastic and endothelial cells via VEGF-C and HGF. Immunol. Cell Biol.

Manaster, I., Mizrahi, S., Goldman-Wohl, D., Sela, H.Y., Stern-Ginossar, N., Lankry, D., Gruda, R., Hurwitz, A., Bdolah, Y., Haimov-Kochman, R., et al. (2008). Endometrial NK Cells Are Special Immature Cells That Await Pregnancy. J. Immunol.

Nancy, P., Tagliani, E., Tay, C.S., Asp, P., Levy, D.E., and Erlebacher, A. (2012). Chemokine gene silencing in decidual stromal cells limits T cell access to the maternal-fetal interface. Science7.

Parham, P., and Moffett, A. (2013). Variable NK cell receptors and their MHC class i ligands in immunity, reproduction and human evolution. Nat. Rev. Immunol.

PrabhuDas, M., Bonney, E., Caron, K., Dey, S., Erlebacher, A., Fazleabas, A., Fisher, S., Golos, T., Matzuk, M., McCune, J.M., et al. (2015). Immune mechanisms at the maternal-fetal interface: Perspectives and challenges. Nat. Immunol.

Seshadri, S., and Sunkara, S.K. (2014). Natural killer cells in female infertility and recurrent miscarriage: A systematic review and meta-analysis. Hum. Reprod. Update.

Sotnikova, N., Voronin, D., Antsiferova, Y., and Bukina, E. (2014). Interaction of decidual CD56+ NK with trophoblast cells during normal pregnancy and recurrent spontaneous abortion at early term of Gestation. Scand. J. Immunol.

Stuart, T., Butler, A., Hoffman, P., Hafemeister, C., Papalexi, E., Mauck, W.M., Hao, Y., Stoeckius, M., Smibert, P., and Satija, R. (2019). Comprehensive Integration of Single-Cell Data. Cell.

Suryawanshi, H., Morozov, P., Straus, A., Sahasrabudhe, N., Max, K.E.A., Garzia, A., Kustagi, M., Tuschl, T., and Williams, Z. (2018). A single-cell survey of the human first-trimester placenta and decidua. Sci. Adv.

Vacca, P., Vitale, C., Montaldo, E., Conte, R., Cantoni, C., Fulcheri, E., Darretta, V., Moretta, L., and Mingari, M.C. (2011). CD34+ hematopoietic precursors are present in human decidua and differentiate into natural killer cells upon interaction with stromal cells. Proc. Natl. Acad. Sci. U. S. A.

Varol, C., Mildner, A., and Jung, S. (2015). Macrophages: Development and Tissue Specialization. Annu. Rev. Immunol.

Vento-Tormo, R., Efremova, M., Botting, R.A., Turco, M.Y., Vento-Tormo, M., Meyer, K.B., Park, J.E., Stephenson, E., Polański, K., Goncalves, A., et al. (2018). Single-cell reconstruction of the early maternal–fetal interface in humans. Nature.

Wallace, A.E., Host, A.J., Whitley, G.S., and Cartwright, J.E. (2013). Decidual natural killer cell interactions with trophoblasts are impaired in pregnancies at increased risk of preeclampsia. Am. J. Pathol.

Wang, F., Zhou, Y., Fu, B., Wu, Y., Zhang, R., Sun, R., Tian, Z., and Wei, H. (2014). Molecular signatures and transcriptional regulatory networks of human immature decidual NK and mature peripheral NK cells. Eur. J. Immunol. 44, 2771–2784.

Wang, W.J., Hao, C.F., Yi-Lin Yin, G.J., Bao, S.H., Qiu, L.H., and Lin, Q. De (2010). Increased prevalence of T helper 17 (Th17) cells in peripheral blood and decidua in unexplained recurrent spontaneous abortion patients. J. Reprod. Immunol.

Wang, Y., Song, F., Zhu, J., Zhang, S., Yang, Y., Chen, T., Tang, B., Dong, L., Ding, N., Zhang, Q., et al. (2017). GSA: Genome Sequence Archive*. Genomics, Proteomics Bioinforma. 15, 14–18.

Wu, X., Jin, L.-P., Yuan, M.-M., Zhu, Y., Wang, M.-Y., and Li, D.-J. (2005). Human First-Trimester Trophoblast Cells Recruit CD56 bright CD16 − NK Cells into Decidua by Way of Expressing and Secreting of CXCL12/Stromal Cell-Derived Factor 1. J. Immunol.

Yang, F., Zheng, Q., and Jin, L. (2019). Dynamic Function and Composition Changes of Immune Cells During Normal and Pathological Pregnancy at the Maternal-Fetal Interface. Front. Immunol.

Yang, H., Qiu, L., Chen, G., Ye, Z., Lü, C., and Lin, Q. (2008). Proportional change of CD4+CD25+ regulatory T cells in decidua and peripheral blood in unexplained recurrent spontaneous abortion patients. Fertil. Steril.

Yu, G. (2015). Statistical analysis and visualization of functional profiles for gene and gene clusters. Bioconductor.Org.

Zhang, J., Dunk, C., Croy, A.B., and Lye, S.J. (2016). To serve and to protect: the role of decidual innate immune cells on human pregnancy. Cell Tissue Res.

Zhang, Z., Zhao, W., Xiao, J., Bao, Y., He, S., Zhang, G., Li, Y., Zhao, G., Chen, R., Gao, Y., et al. (2020). Database Resources of the National Genomics Data Center in 2020. Nucleic Acids Res.

Zheng, G.X.Y., Terry, J.M., Belgrader, P., Ryvkin, P., Bent, Z.W., Wilson, R., Ziraldo, S.B., Wheeler, T.D., McDermott, G.P., Zhu, J., et al. (2017). Massively parallel digital transcriptional profiling of single cells. Nat. Commun.

Zhou, Y., Fu, B., Xu, X., Zhang, J., Tong, X., Wang, Y., Dong, Z., Zhang, X., Shen, N., Zhai, Y., et al. (2020). PBX1 expression in uterine natural killer cells drives fetal growth. Sci. Transl. Med.

